# Competitive multiplication rate variation of *Plasmodium falciparum* isolates is determined by differences in intrinsic potential

**DOI:** 10.64898/2026.08.04.742750

**Authors:** Lindsay B. Stewart, James Philpott, Gordon A. Awandare, David J. Conway

## Abstract

Malaria parasite virulence will be impacted by naturally occurring variation in multiplication rates. Clinical isolates of *Plasmodium falciparum* exhibit a range of multiplication rates under exponential growth conditions in culture, but it needs to be discovered if relative multiplication rates under competitive conditions are principally defined by this underlying multiplication rate potential. Relative multiplication rates of *P. falciparum* lines were investigated in 14-day competition assays against a standard competitor clone HB3, with other long-term laboratory-adapted clones showing similar or higher mean per-48-hour rates ranging from 0.98 to 1.51 relative to HB3. In contrast, thirteen Ghanaian clinical isolates cultured for close to two months prior to assay had lower rates, ranging from 0.46 to 0.76 relative to HB3. Analysing the single-genotype clinical isolates and laboratory-adapted clones, there was a highly significant correlation between the competitive rates and previously determined exponential rates (Spearman’s rho = 0.95, P = 0.0003), which accounted for most of the observed variance (Pearson’s *r*^2^ = 0.77, P = 0.004). Testing the effect of alternative nutritional supplementation with human serum or Albumax showed a majority of parasites had consistent competitive rates under either condition, indicating that intrinsic differences determine most variation among parasites.

## Introduction

The malaria parasite *Plasmodium falciparum* causes more human deaths than any other eukaryotic pathogen. Every year, approximately 600,000 deaths and more than 200 million clinical infections are caused by blood-stage parasites replicating to high densities ^1,2^. However, many infections are asymptomatic with parasites maintained in the blood at levels that are undetectable or that can only be detected by sensitive PCR tests ^3^, providing an endemic reservoir that is a major challenge for malaria elimination. Low parasite densities are not only due to suppression by acquired host immunity, but are relatively common in individuals without any prior immunity in areas of low endemicity, suggesting the possibility of intrinsically low multiplication rates ^4^.

There is a wide range of evidence that *P. falciparum* multiplication rates vary. Laboratory-adapted strains show differences in their multiplication rates in culture ^5,6^, although such differences might be affected by laboratory-acquired mutations ^7–10^. Studies of laboratory-adapted parasites in artificially induced human infections have indicated multiplication rate differences between strains, although data are limited and vary between study sites ^11–15^. Investigations on clinical isolates in culture have shown multiplication rate variation, either in the initial *ex vivo* cycle ^16–18^, or during early continuous cultivation before mutants become common ^5,9^. These latter studies show that parasite multiplication rates in clinical isolates under exponential growth conditions range from less than 2-fold to approximately 8-fold per 48-hour period (corresponding to a typical intra-erythrocytic cycle) ^5,9^. Significantly, positive correlations have been shown between exponential multiplication rates in culture and parasitaemia levels of the sampled infections from which isolates were derived ^9,19^.

Clearly, exponential growth assays need to exclude variations in conditions that parasites face during infections. Parasites naturally face a wide range of conditions that include resource constraints and competition with each other, particularly when parasite densities are high. Competitive co-culture assays have been used to test for differences in relative growth rates between different *P. falciparum* cultured lines including unrelated isolates ^20^, sibling progeny of experimental crosses ^21,22^, or genetically engineered parasites ^23–25^. Although it is not feasible to test many parasite lines in all possible combinations, pairwise tests among a small number of clones have indicated that a rank order may be derived without conducting all comparisons ^20^. This suggests that the relative rates of a larger number of parasite lines may be effectively determined by comparisons against a standard competitor clone. However, one study indicated that competition between clones differed depending on culture medium supplementation ^21^, suggesting variation in nutritional requirements which needs further investigation.

It is important to determine whether relative multiplication rates of parasites under competitive conditions are mainly defined by the underlying exponential multiplication rate potential of each isolate, or other modifying effects operate. Here, relative multiplication rates of long-term laboratory-adapted *P. falciparum* clones and short-term cultured clinical isolates were investigated in co-cultures with a standard competitor clone. This showed a wide range of competitive rates, with the long-term laboratory-adapted clones having similar or higher rates than the standard competitor, while all of the clinical isolates had lower rates. Notably, there was a highly significant correlation between the competitive rates and the exponential rates previously determined, which explained most of the variance. Comparing alternative medium supplementation with human serum or serum-free nutrients showed minor effects, with most parasites showing consistent relative rates, indicating that variation in these rates is mostly determined by intrinsic differences between parasites.

## Results

### Paired co-culture of different *P. falciparum* lines with allele-specific discrimination

Four laboratory-adapted *P. falciparum* laboratory clones and thirteen clinical isolates were analysed, each of which had exponential multiplication rates previously determined ^5,6,9^ (Table 1). As the laboratory-adapted clone HB3 could be discriminated from each of the other individual clones and isolates by allele-specific qPCR methods, this was selected as a standard competitive clone in co-culture. It was previously indicated that HB3 has an exponential multiplication rate of approximately 8-fold per 48 hours, similar to 3D7 and D10, and slower than Dd2 which multiplied at approximately 10-fold (Table 1) ^5^. Thirteen Ghanaian clinical isolates were also selected for testing relative growth rates in co-culture with clone HB3. The exponential multiplication rates of each of these individual isolates after 25 days of culture were previously reported, ranging from 2.5-fold to 6.9-fold per 48 hours (Table 1).

**Table 1.**
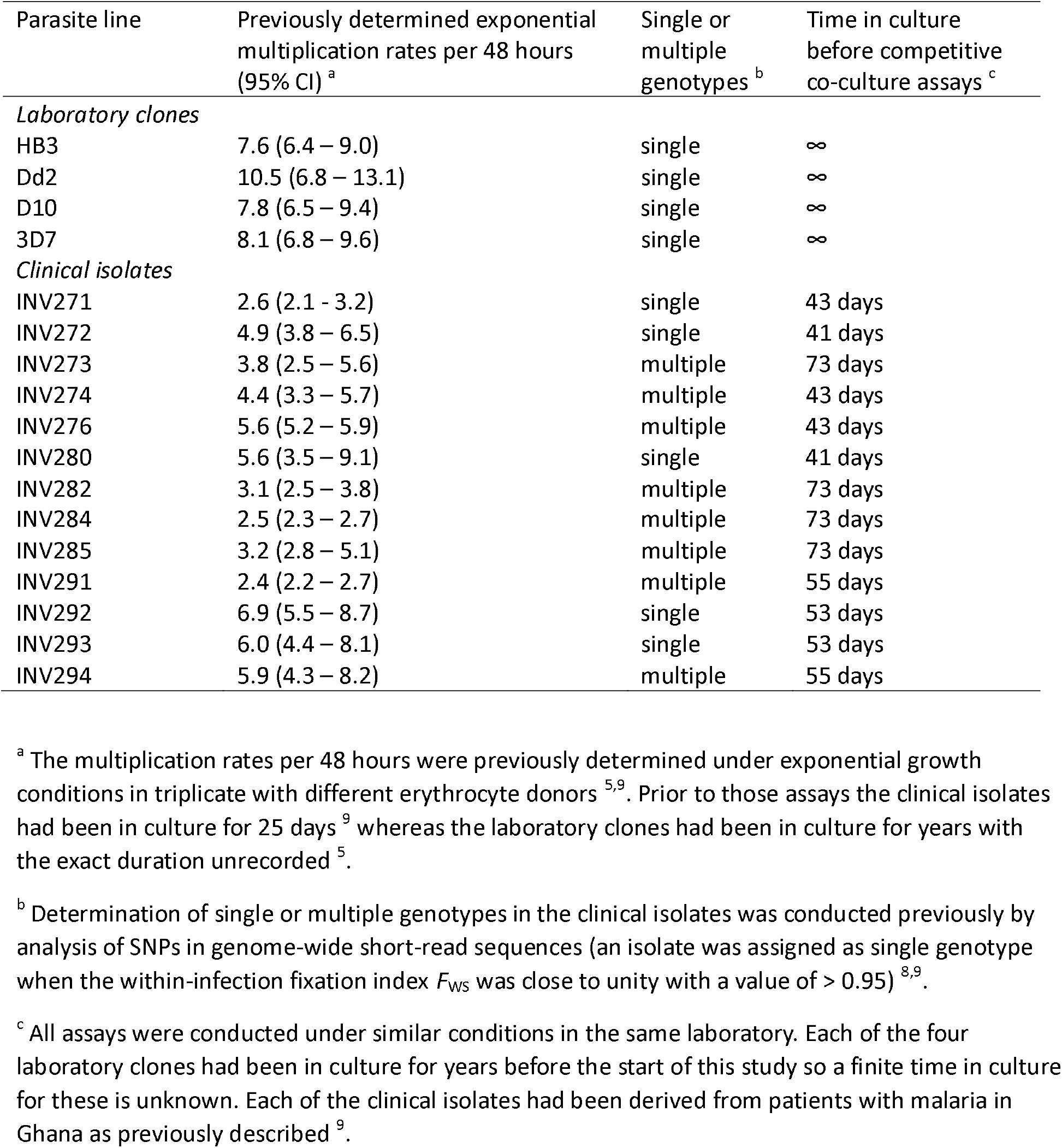
*P. falciparum* laboratory clones and clinical isolates selected for analysis in competitive multiplication rate assays.

Each parasite line was tested in competition with clone HB3 over a 14-day period using triplicate co-cultures. The relative multiplication rate in each replicate was expressed as an overall fold difference in rate per 48 hours compared to HB3, and the assay result for each line was expressed as the mean and standard deviation across the replicates (Figure 1).

**Figure 1.**
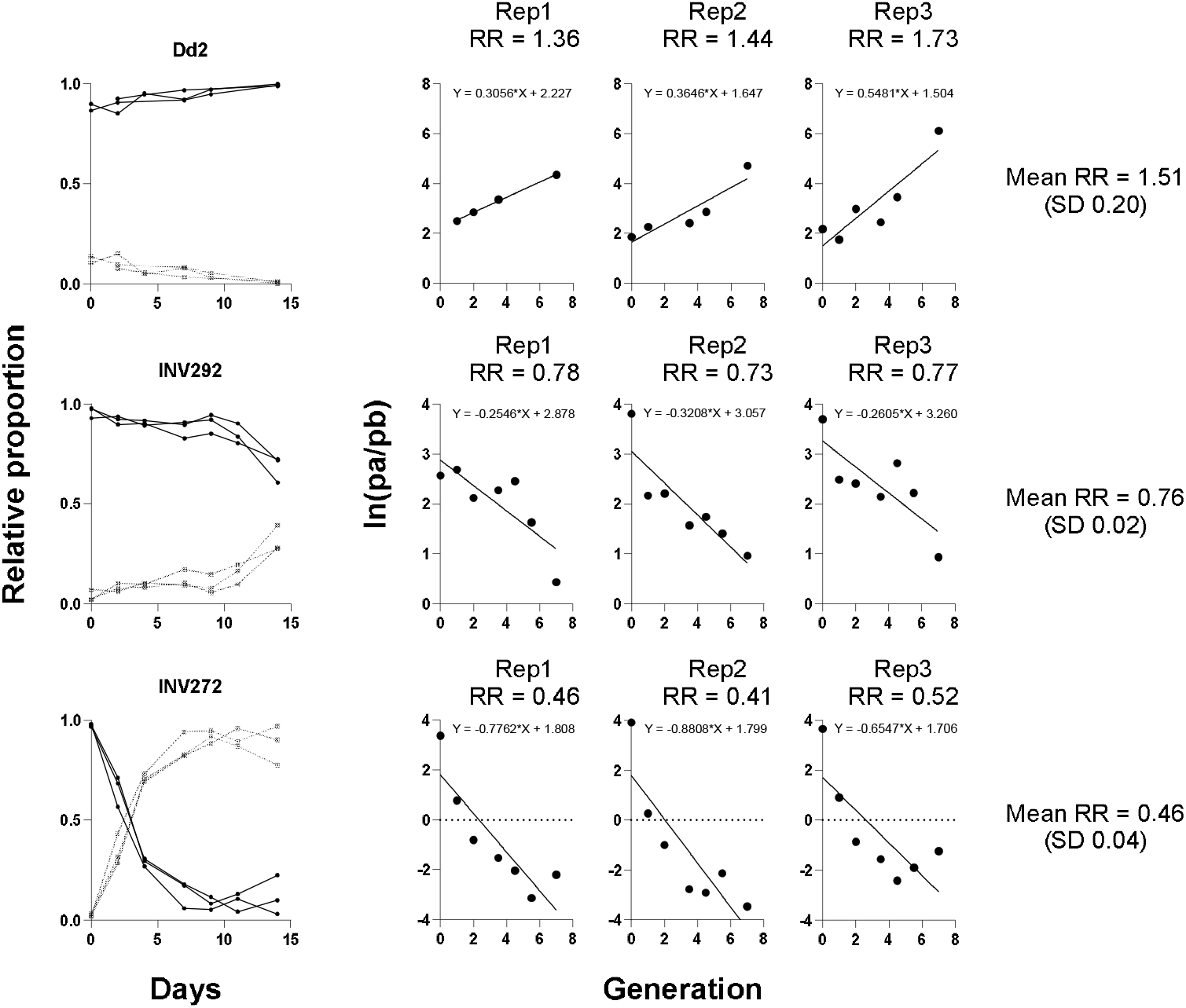
Relative multiplication rate co-culture assays of different *P. falciparum* lines tested in competition with clone HB3. Examples are shown of an assay of clone Dd2 which multiplies faster than HB3, and of two Ghanaian clinical isolates (INV272 and INV292) which multiply slower than HB3. Assays were conducted over a 14-day period with triplicate co-cultures in different donor erythrocytes. The left panel for each parasite line show the proportions in the triplicate co-cultures over time, with each tested parasite line in black and HB3 as the shadow lines in grey (HB3 was intentionally started as a minority proportion in all assays). The triplicate panels to the right show the natural log of the ratio of proportions of each tested parasite line versus HB3 [ln ratio (a/b)] plotted to determine overall relative rate of change over time. The relative multiplication rate in each of the replicates was expressed as an overall fold difference in rate per 48 hours compared to HB3, and the assay result for each line is expressed as the mean and standard deviation across the replicates. In these plots, 48-hour periods are labelled as ‘generations’ corresponding to a nominally typical parasite cycle length in order to derive a standardised relative rate fold-difference compared to HB3. The mean and standard deviation of the relative rate compared to HB3 for each assay is shown on the right. The numerical data for these assays are given in Supplementary Table 1 and the Supplementary Dataset.

### Competitive co-culture of laboratory-adapted *P. falciparum* clones

To assess differences in their relative multiplication rates, each of the unrelated laboratory-adapted parasite clones was tested in triplicate 14-day co-cultures with clone HB3 on two or three different occasions. The fastest growing clone was Dd2, with relative per-48-hour multiplication rates on three different occasions being 1.51, 1.42, and 1.56-fold higher than HB3 (Figure 2 and Table 2). Clone D10 was also consistently faster growing than HB3 in multiple replicate assays on two different occasions by 1.32 and 1.45-fold. Clone 3D7 showed growth that was similar to that of HB3 on both occasions, 0.97 and 0.98-fold (Figure 2 and Table 2).

**Table 2.** Relative multiplication rates of *P. falciparum* laboratory clones and clinical isolates compared to standard clone HB3 in competitive co-culture assays.

| Parasite line | Relative Rate versus HB3 per 48 hours (from 14-day competition assays) |  |  |  |
| --- | --- | --- | --- | --- |
|  | Replicate 1 | Replicate 2 | Replicate 3 | Mean (SD) |
| *Laboratory clones* |  |  |  |  |
| Dd2 - assay 1 | 1.36 | 1.44 | 1.73 | 1.51 (0.20) |
| Dd2 - assay 2 | 1.50 | 1.38 | 1.41 | 1.43 (0.05) |
| Dd2 - assay 3 | 1.50 | 1.59 | 1.59 | 1.56 (0.04) |
| D10 - assay 1 | 1.48 | 1.26 | 1.31 | 1.35 (0.09) |
| D10 - assay 2 | 1.43 | 1.47 | 1.45 | 1.45 (0.02) |
| 3D7 - assay 1 | 0.99 | 0.94 | 0.97 | 0.97 (0.02) |
| 3D7 - assay 2 | 0.89 | 1.10 | 0.94 | 0.98 (0.09) |
| *Clinical isolates* |  |  |  |  |
| INV271 | 0.57 | 0.53 | 0.53 | 0.55 (0.02) |
| INV 272 | 0.46 | 0.41 | 0.52 | 0.46 (0.05) |
| INV 273 | 0.79 | 0.74 | 0.71 | 0.75 (0.03) |
| INV 274 | 0.63 | 0.63 | 0.53 | 0.60 (0.05) |
| INV 276 | 0.65 | 0.53 | 0.63 | 0.60 (0.05) |
| INV 280 | 0.65 | 0.68 | 0.61 | 0.65 (0.03) |
| INV 282 | 0.62 | 0.68 | 0.63 | 0.64 (0.02) |
| INV 284 | 0.63 | 0.67 | - | 0.65 (0.03) |
| INV 285 | 0.66 | 0.74 | 0.69 | 0.70 (0.03) |
| INV 291 | 0.69 | 0.65 | 0.66 | 0.67 (0.01) |
| INV 292 | 0.78 | 0.73 | 0.77 | 0.76 (0.03) |
| INV 293 | 0.68 | 0.63 | 0.66 | 0.65 (0.03) |
| INV 294 | 0.67 | 0.61 | 0.57 | 0.62 (0.04) |

**Figure 2.**
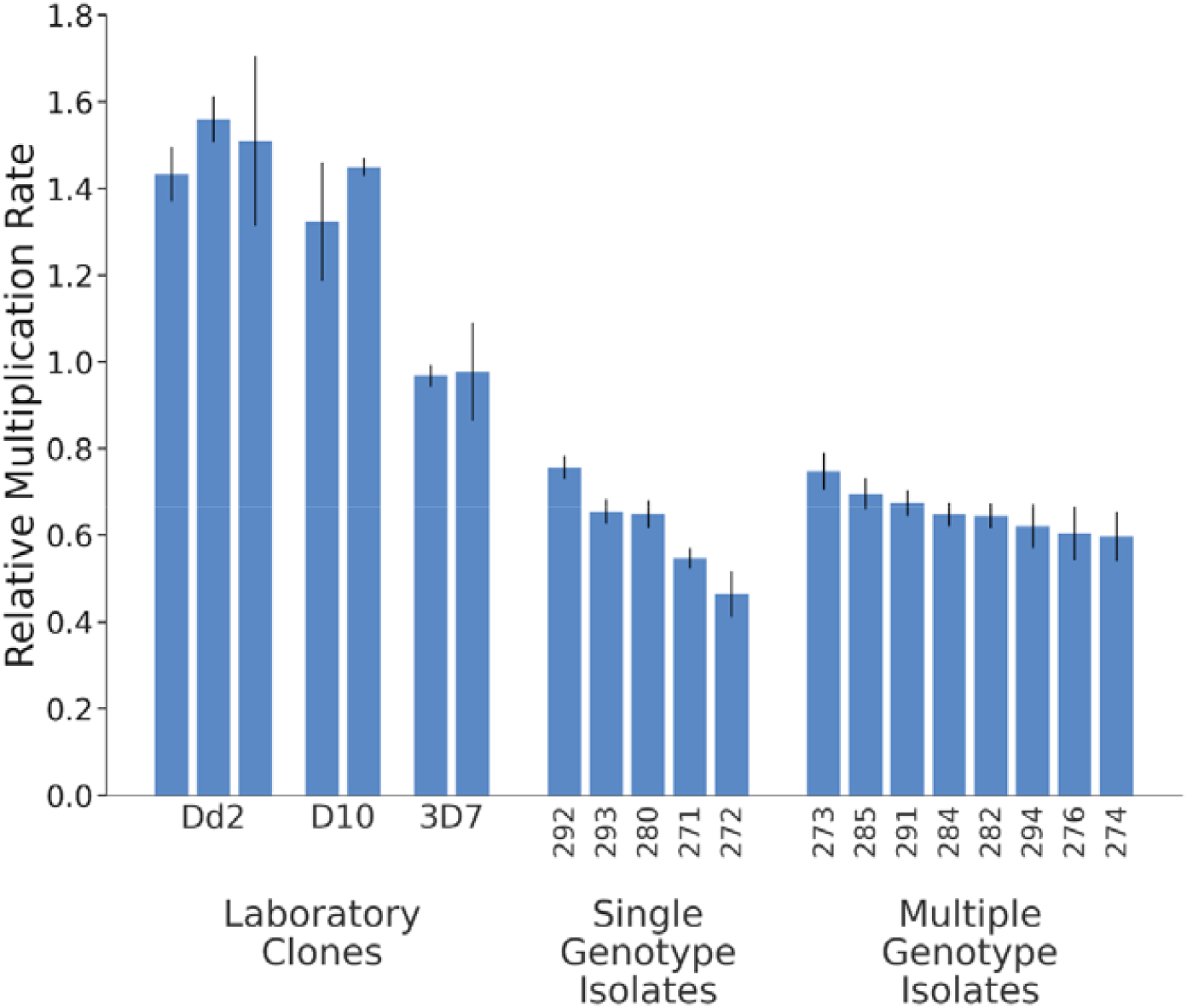
Relative multiplication rates of different *P. falciparum* laboratory clones and clinical isolates in competition with clone HB3. Each assay involved triplicate co-cultures with clone HB3 over 14 days, with erythrocytes from different donors in each replicate (as illustrated for two of the parasite lines in Figure 1). Bars show the mean relative rate determined for each parasite line in each assay compared to HB3 having a reference value of 1.0 and expressed as overall per 48-hour relative rate). Vertical lines show standard deviations of the means across replicates. Laboratory-adapted clones were each assayed on more than one occasion (three times for clone Dd2 and twice for the others) with consistent results across assays. The clinical isolates were each assayed on one occasion, after having been in culture for a mean of 55 days (Table 1). These isolates are identified by their numbers in the ‘INV’ series listed in Table 1. Previous genome sequencing indicated that five of these isolates contained single genotypes and the other eight contained multiple genotypes^8^. Numerical results of each of the assays are given in Table 2, with all data on each replicate given in Supplementary Table 1 and the Supplementary Dataset.

The similarity of the multiplication rate of 3D7 to HB3 under competitive conditions accords with these clones having similar multiplication rates under exponential conditions (Table 1). Likewise, clone Dd2 had a faster multiplication rate than HB3 when previously tested under exponential conditions, as well as a faster rate in all of the direct competition assays. However, clone D10 had a faster rate in competition with HB3 although their previously determined exponential rates were similar.

### Competitive co-culture of Ghanaian clinical isolates paired with standard clone HB3

Thirteen cultured clinical isolates previously analysed for exponential multiplication rate variation ^9^ were selected for testing in competitive co-culture with clone HB3. These had been cultured for a mean of 55 days (range 41 to 73 days) prior to the assays, and five of them had single genotypes while eight contained multiple genotypes (Table 1). The clinical isolates showed a range of multiplication rates from 0.46 to 0.76, as mean per-48-hour relative rates compared with HB3. These rates were similar for single genotype and multiple genotype isolates, with a slightly narrower range among the multiple genotype isolates (Figure 2, Supplementary Table 1).

### Correlation between competitive and exponential multiplication rates

Across all of the parasite lines tested, including the mixed-genotype isolates, there was a significant positive rank correlation between the competitive multiplication rates relative to HB3 and the exponential multiplication rates previously determined (Spearman’s rho = 0.518, P = 0.04). However, exponential multiplication rates of the clinical isolates had been assayed after 25 days of culture, and previous analysis showed that there were subsequent changes in genomic composition of the mixed-genotype isolates in culture ^8^, so they would not be equivalent when competitive assays were conducted (after between 41 and 73 days in culture as noted in Table 1). Excluding these mixed-genotype isolates, there was a much stronger rank correlation between the competitive multiplication rates and exponential multiplication rates (Spearman’s rho = 0.952, P = 0.0003), and also a strong parametric correlation (Pearson’s r = 0.877, P = 0.004) (Figure 3).

**Figure 3.**
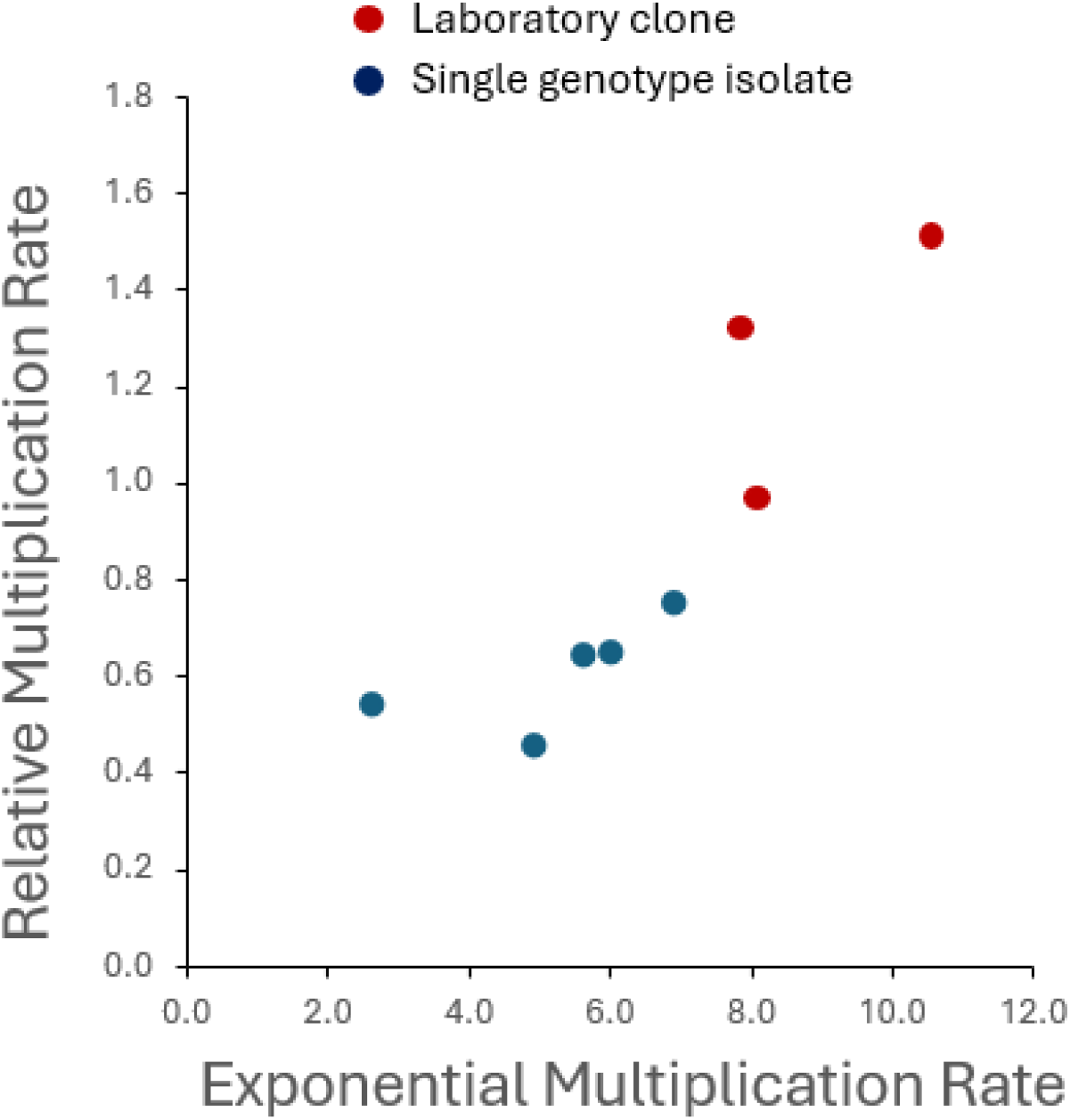
Correlation between relative multiplication rates assayed by competitive co-culture and exponential multiplication rates determined in separate assays. The relative rates were determined in the present study by 14-day co-culture with the standard competitor clone HB3 (Table 2), and the exponential multiplication rates were previously determined (Table 1). The analysis focuses on clones and single genotype isolates, with mixed genotype isolates being excluded. The competitive rates were highly correlated with the independently measured individual exponential rates (Spearman’s rho = 0.95, P = 0.0003; Pearson’s r = 0.88, P = 0.004), with the square of the parametric correlation coefficient suggesting approximately three-quarters of the variance was explained (Pearson’s *r* = 0.77).

The square of the parametric correlation coefficient suggested approximately three-quarters of the variance in relative rates under competitive conditions may be explained by the underlying exponential multiplication rate potential of each parasite line (Pearson’s *r*^2^ = 0.77)(Figure 3). However, the scatter of points in the correlation suggests that additional effects may operate to influence the competitive rates of some isolates.

### Testing for variation under different nutritional conditions

It was investigated whether parasite clones vary in their relative multiplication rates depending on the type of nutritional supplementation provided. Relative rates of eight clones were tested in competitive co-culture with HB3 in media supplemented with either Albumax or serum. In addition to long-term laboratory adapted clones, six of the clones tested were more recently derived from Ghanaian isolates ^8^, one clone being selected for testing from each of six parental isolates so that each was unrelated.

The competitive multiplication rates of most clones compared to HB3 were similar under either condition (Figure 4). A few clones showed minor differences between the conditions, although none showed a reversal relative to HB3. An example of a clone showing a trend in each direction was noted, as clone Dd2 had more of a competitive advantage in Albumax than in human serum while clone 286_B10 had more of a disadvantage in Albumax (Figure 4, Supplementary Table 2 and Supplementary Dataset). The observation that most clones had consistent relative rates, with only minor differences between conditions, supports an interpretation that intrinsic parasite differences predictably determine most of the variation in competition.

**Figure 4.**
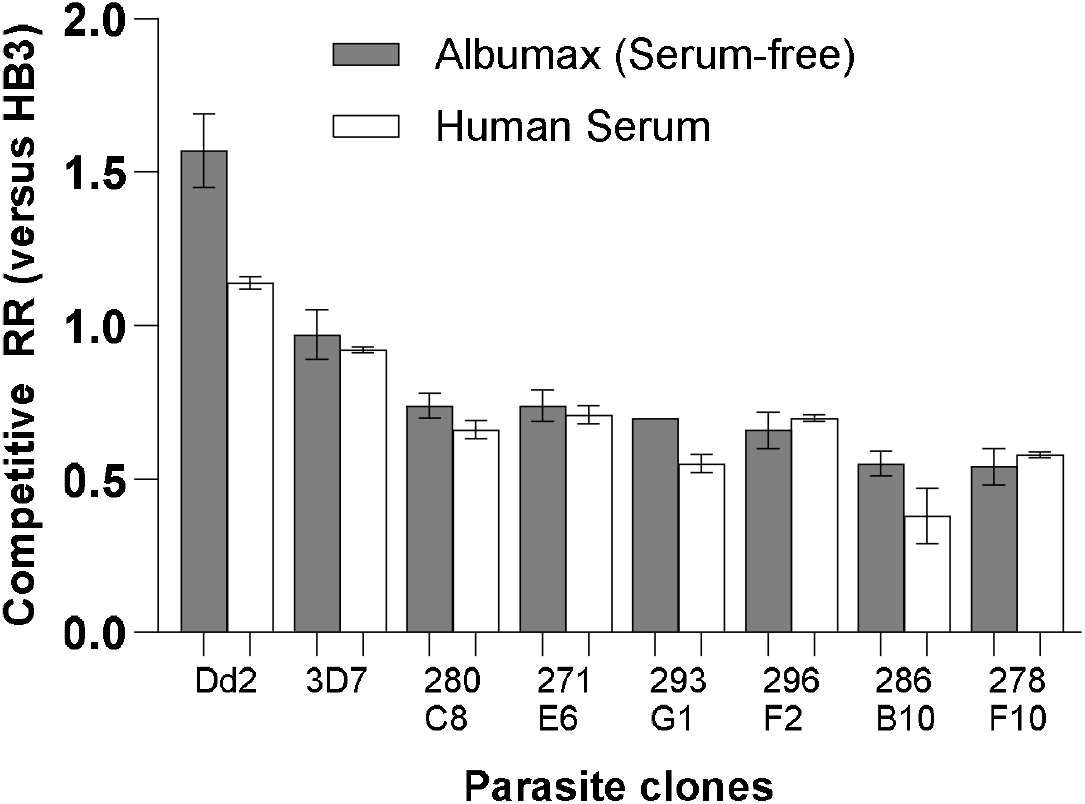
Comparison of the effects of culture supplementation with either serum or Albumax on the relative multiplication rates of different clones in co-culture with the standard HB3 competitor clone. The competition assays were performed over 14 days as described for the previous assays. And each comparison was performed in duplicate or triplicate. Relative rates (RR) are expressed as overall mean per-48 hour differences compared to clone HB3 having a reference value of 1.0. Most of the clones had similar relative rates under either condition, and none showed a competitive reversal relative to HB3. Particular clones showed minor differences, with one example in each direction highlighted (Dd2 had more of a competitive advantage versus HB3 in Albumax than in human serum while clone 286_B10 had more of a disadvantage in Albumax than in human serum). Numerical results for all replicates are given in Supplementary Table 2 (full assay data are given in the Supplementary Dataset).

## Discussion

This study shows an overall strong correlation between the relative multiplication rates of different *P. falciparum* lines in competitive co-culture conditions and the intrinsic rates of each line assayed separately under exponential conditions. This indicates that a large part of the variance in competitive rates is explainable by underlying exponential rate potential, while other determinants operating under non-exponential growth conditions are likely to contribute to the observed residual variation.

It remains to be determined which phases of the developmental cycle contribute most to this variation ^26^. Artificial experimental disruption of individual genes affects growth efficiency of long-term culture-adapted parasites ^10,27^, and some of these mutants influence tolerance of heat shock or oxidative stress ^28^. Naturally occurring multiplication rate variation might involve some of those genes, or may be largely regulated by others. Although the replication cycle of *P. falciparum* involves co-ordinated transcriptional networks ^29–32^, it is not yet known if these networks control the multiplication rate variation among different isolates.

This study utilised a single competitor clone in all of the assays, so that the relative multiplication rates of diverse parasite lines were compared to this as a standard reference. The clone HB3 was utilised as it had previously shown an exponential multiplication rate of approximately 8-fold per 48 hours ^5^, similar to that of other laboratory-adapted clones, although less than that of clone Dd2 which has consistently been shown to have a higher rate ^5,6^. Another useful feature of clone HB3 is that it has a relatively consistent gametocyte conversion rate of approximately 10% per cycle in repeated tests from continuous cultures, whereas some other laboratory adapted clones including Dd2 and NF54 have more temporally varying rates ^33^. It is generally recognised that variation in sexual commitment rates of different parasite lines will affect their multiplication rates in culture. As most clinical isolates have sexual commitment rates ranging from approximately 3 % to 15 % per cycle this would contribute marginally to the variation seen. Future work on assays of parasites in co-culture could consider use of a genetically modified competitor clone that does not undergo any sexual commitment.

Although the underlying exponential multiplication rate potential explained most of the competitive variation, it was considered that parasites could vary in their responses to the co-culture conditions that differ from those supporting exponential growth. Considering variations that parasites will experience in natural infections, nutritional components are likely to be important. Previous investigation of competitive growth indicated alternative nutritional medium supplements may favour parasites differentially, and a separate comparison of two different laboratory-adapted clones indicated that parasite gene expression is significantly affected in culture with Albumax compared with human serum supplementation ^34^. Therefore, the effect of human serum versus Albumax supplementation was tested on eight genotypically different clones here, showing that most had consistent relative rates, with only minor differences seen between conditions. This indicates that intrinsic parasite differences reproducibly determine most of the competitive variation.

It is possible that the transcriptional profile of each parasite clone may be variable, with different expression types represented in different individual parasites. Single-cell RNA sequencing of isolates under competitive growth conditions could enable pseudotime analysis to estimate variation in cycle development time, and the developmental staging of individual parasites ^35^. Such data would also reveal whether a variable minority of early trophozoite-stage parasites undergo reversible-arrested development, hypothesised as an adaptive mechanism of escaping metabolic or drug-induced stress *in vivo* ^36^.

To identify targets for future therapeutics and vaccines, it is vital to discover the causes of intrinsic variation in malaria parasite multiplication rates. The use of competitive assays enables the differences in multiplication rates among parasite lines to be investigated under a wider range of conditions than those that allow exponential multiplication. Further application of the competitive co-culture assays will enable testing whether parasites vary in dependence on particular nutrients, or on properties of host erythrocytes, or might respond to modifications of the environment by other parasites.

## Methods

### *P. falciparum* laboratory clones and cultured clinical isolates

Four long-term laboratory-adapted *P. falciparum* clones (HB3, Dd2, D10, and 3D7) and 13 cultured *P. falciparum* clinical isolates were studied (Supplementary Table 1). The clinical isolates were selected from among those previously assayed under exponential growth conditions ^9^. These isolates were originally sampled from uncomplicated clinical malaria patients in Ghana with approval by the Ethics committees of the Ghana Health Service, the Noguchi Memorial Institute for Medical Research at the University of Ghana, the Navrongo Health Research Centre and the London School of Hygiene and Tropical Medicine. Patients were aged 2 – 14 years, and written informed consent was obtained from parents or other legal guardians of all participants. All methods were performed in accordance with the relevant guidelines and regulations.

The *P. falciparum* clinical isolates had been in culture for a mean of 53 days (range from 41 to 73 days) before assay for relative multiplication rates in this study in co-culture with P. clone HB3. In addition, six previously described *P. falciparum* clones derived from Ghanaian isolates ^8^ were used to compare relative multiplication rates in co-culture with clone HB3 under alternative nutritional supplementation with either Albumax or human serum.

### Malaria parasite culture conditions

*P. falciparum* laboratory-adapted clones and short-term cultured clinical isolates were thawed from glycerolyte preservation and cultured at 37°C using standard methods. Briefly, 12% NaCl was added dropwise to each cryopreserved sample (adding the equivalent of half of the volume of the cryopreserved material) while gently agitating the tube to allow mixture. Following this, the tube was left to stand for 5 min, then 1.6% NaCl was added dropwise (adding 10 times the original volume of the cryopreserved material), gently agitating to allow mixture. After centrifugation for 5 min at 500 g, the supernatant was removed and cells were resuspended in the same volume of RPMI 1640 medium (Sigma-Aldrich, UK) containing 0.5% Albumax™ II (Thermo Fisher Scientific, UK). Cells were centrifuged again, supernatant removed and the erythrocyte pellet (comprising at least 250 µl for each sample) was resuspended at 3% haematocrit in RPMI 1640 medium supplemented with 0.5% Albumax II, under an atmosphere of 5% O_2_, 5% CO_2_, and 90% N_2_, with orbital shaking of flasks at 60 revolutions per minute.

### Allele-specific qPCR assays to discriminate different *P. falciparum* cultured lines

Published methods were used for quantitative discrimination of different *P. falciparum* clones and isolates in co-culture, based on allele-specific qPCR assays ^5,6,37^. These assays allow quantitative assessment of relative proportions of different *P. falciparum* genotypes, by discriminating alleles of antigen genes with divergent sequence regions, a trimorphic block 2 region of *msp1* ^37^, a dimorphic block 4 region of *msp1* ^6^, and a dimorphic region of *msp6* ^5^. Previous application of these assays in this laboratory have confirmed their precision in enabling quantitative assessment of the ratios of different parasite clones in co-culture over time ^5^. Consideration of the genotypes of parasites in the current study showed that clone HB3 could be discriminated from all other clones and isolates at one or more of the loci, so that it could serve as a standard competitor for quantitative assessment of relative multiplication rates. Conveniently, HB3 could be discriminated from all other clones and isolates using a single protocol of allele-specific qPCR targeting block 4 of the msp1 gene ^6^ (only HB3 had the *HB3-like* allele while all others had the *Dd2-like* allele), so this was used for all assays.

### Competitive co-culture assays to determine relative multiplication rates

For each competition assay, fresh blood was procured (commercially from Cambridge Bioscience, UK) from three anonymous blood donors in the UK who had not recently taken any antimalarial drugs or travelled to a malaria endemic area, and who did not have sickle-cell trait or other known haemoglobin variants. The erythrocytes were stored at 4°C for no more than 2 days then washed immediately before use in the assays. Each assay was performed in triplicate, with erythrocytes from different donors in separate flasks or wells of 6-well plates.

Paired co-culture assays were initiated using seeding cultures of clone HB3 and each competitor diluted into a starting ratio of approximately 1:10, with an overall parasitaemia of approximately 0.2%. Given that HB3 has an exponential multiplication rate of approximately 8-fold per 48 hours ^5^, and most clinical isolates (including the Ghanaian isolates analysed here) have lower exponential rates ^5,9,19,38^, the assay was designed to have HB3 starting as a minority proportion so that a trajectory of change in proportions would be effectively sampled over 14 days. Each replicate was performed in 10 ml volume culture in a flask or 5 ml volume in a 6-well plate, at a haematocrit of 3%.

A 300 µl volume of suspended culture was sampled from each replicate every 3 or 4 days for DNA extraction and qPCR, and each replicate culture was diluted by approximately 1 in 20 with additional erythrocytes from the same donor. Numbers of copies of each of the alternative competing genotypes were measured by qPCR and expressed as a proportion of the total copies observed. The natural log of the ratio of proportions of each tested parasite line versus HB3 [ln ratio (a/b)] is plotted to determine overall rate of change over time, as the slope of the line (*s*) for each replicate over the 14 days. The anti-log value of s yields the multiplication rate ratio (RR) relative to HB3 for each culture replicate. The mean and standard deviation of RR across triplicate co-cultures is expressed as the individual assay result for each parasite isolate or clone.

### Comparisons of parasite competitiveness with different nutrient supplementation

Relative multiplication rates of selected *P. falciparum* clones in competition with HB3 were tested in under alternative media supplement conditions. Paired comparisons were conducted for each replicate in either duplicate or triplicate assays, supplemented with either 0.5% Albumax II or 10% human serum pooled from blood group AB donors (PAN-Biotech). The assay conditions were as described above, with each replicate being conducted in a 5 ml volume on a 6-well plate. In addition to the long-term laboratory adapted clones, six of the clones tested here (271-E6, 278-F10, 280-C8, 286-B10, 293-G1 and 296-F2) were more recently derived from Ghanaian clinical isolates (each clone from a different isolate so that they were unrelated) ^8^.

## Data availability

All data supporting the findings of this study are available within the paper and its Supplementary Information.

