## Supplementary Table for "Competitive multiplication rate variation of *Plasmodium falciparum* isolates is determined by differences in intrinsic potential"

**Supplementary Information**

**Supplementary Table 1.** Relative rates of different *P. falciparum* cultured lines (laboratory adapted clones and clinical isolates) in competition with clone HB3

| Assay | Parasite line | Natural log slope from each replicate [a] | | | Relative rate compared to HB3 in each replicate | | | Mean | SD |
|---|---|---|---|---|---|---|---|---|---|
| | | Rep1 (ln) | Rep2 (ln) | Rep3 (ln) | Rep1 (anti-ln) | Rep2 (anti-ln) | Rep3 (anti-ln) | | |
| 1 | Dd2 | 0.31 | 0.36 | 0.55 | 1.36 | 1.44 | 1.73 | **1.51** | **0.20** |
| 1 | 3D7 | -0.01 | -0.06 | -0.03 | 0.99 | 0.94 | 0.97 | **0.97** | **0.02** |
| 1 | D10 | 0.39 | 0.23 | 0.27 | 1.48 | 1.26 | 1.31 | **1.35** | **0.09** |
| 1 | 271 | -0.56 | -0.63 | -0.63 | 0.57 | 0.53 | 0.53 | **0.55** | **0.02** |
| 1 | 274 | -0.47 | -0.45 | -0.63 | 0.63 | 0.63 | 0.53 | **0.60** | **0.05** |
| 1 | 276 | -0.43 | -0.63 | -0.47 | 0.65 | 0.53 | 0.63 | **0.60** | **0.05** |
| 1 | 291 | -0.38 | -0.43 | -0.41 | 0.69 | 0.65 | 0.66 | **0.67** | **0.01** |
| 1 | 294 | -0.40 | -0.49 | -0.55 | 0.67 | 0.61 | 0.57 | **0.62** | **0.04** |
| 2 | Dd2 | 0.41 | 0.32 | 0.35 | 1.50 | 1.38 | 1.41 | **1.43** | **0.05** |
| 2 | 3D7 | -0.12 | 0.10 | -0.06 | 0.89 | 1.10 | 0.94 | **0.98** | **0.09** |
| 2 | D10 | 0.36 | 0.38 | 0.37 | 1.43 | 1.47 | 1.45 | **1.45** | **0.02** |
| 2 | 285 | -0.41 | -0.31 | -0.37 | 0.66 | 0.74 | 0.69 | **0.70** | **0.03** |
| 2 | 282 | -0.47 | -0.39 | -0.46 | 0.62 | 0.68 | 0.63 | **0.64** | **0.02** |
| 2 | 273 | -0.23 | -0.30 | -0.34 | 0.79 | 0.74 | 0.71 | **0.75** | **0.03** |
| 2 | 284 | -0.46 | -0.41 | - | 0.63 | 0.67 | - | **0.65** | **0.02** |
| 3 | Dd2 | 0.40 | 0.46 | 0.47 | 1.50 | 1.59 | 1.59 | **1.56** | **0.04** |
| 3 | 272 | -0.78 | -0.88 | -0.65 | 0.46 | 0.41 | 0.52 | **0.46** | **0.04** |
| 3 | 280 | -0.44 | -0.39 | -0.49 | 0.65 | 0.68 | 0.61 | **0.65** | **0.03** |
| 3 | 292 | -0.25 | -0.32 | -0.26 | 0.78 | 0.73 | 0.77 | **0.76** | **0.02** |
| 3 | 293 | -0.39 | -0.47 | -0.42 | 0.68 | 0.63 | 0.66 | **0.65** | **0.02** |

[a] The data for each assay including the slopes of the natural logs of proportions over each 14-day assay are given in the Supplementary Dataset.

Assays 1 and 3 were conducted in flasks, Assay 2 was conducted in 6-well plates.

**Supplementary Table 2.** Relative rates of different clones in competition with clone HB3 in different conditions of medium supplementation [a]

| Clone and condition [a] | Rep1 (ln) | Rep2 (ln) | Rep3 (ln) | Rep1 (anti-ln) | Rep2 (anti-ln) | Rep3 (anti-ln) | Mean | SD |
|---|---|---|---|---|---|---|---|---|
| **Dd2 A** | 0.52 | 0.37 | - | 1.69 | 1.45 | - | **1.57** | **0.12** |
| **Dd2 S** | 0.11 | 0.15 | - | 1.12 | 1.16 | - | **1.14** | **0.02** |
| **3D7 A** | 0.05 | -0.12 | - | 1.06 | 0.89 | - | **0.97** | **0.08** |
| **3D7 S** | -0.09 | -0.08 | - | 0.92 | 0.93 | - | **0.92** | **0.01** |
| **271_E6 A** | -0.36 | -0.33 | -0.21 | 0.70 | 0.72 | 0.81 | **0.74** | **0.05** |
| **271_E6 S** | -0.30 | -0.32 | -0.39 | 0.74 | 0.72 | 0.68 | **0.71** | **0.03** |
| **278_F10 A** | - | -0.74 | -0.52 | - | 048 | 0.59 | **0.54** | **0.06** |
| **278_F10 S** | - | -0.51 | -0.60 | - | 0.60 | 0.55 | **0.58** | **0.01** |
| **280_C8 A** | -0.26 | -0.35 | - | 0.77 | 0.70 | - | **0.74** | **0.04** |
| **280_C8 S** | -0.45 | -0.37 | - | 0.64 | 0.69 | - | **0.66** | **0.03** |
| **286_B10 A** | -0.66 | -0.51 | -0.66 | 0.52 | 0.60 | 0.52 | **0.55** | **0.04** |
| **286_B10 S** | -1.34 | -0.76 | -0.91 | 0.26 | 0.47 | 0.40 | **0.38** | **0.09** |
| **293_G1 A** | -0.35 | -0.35 | - | 0.70 | 0.70 | - | **0.70** | **0.00** |
| **293_G1 S** | -0.65 | -0.53 | - | 0.52 | 0.59 | - | **0.55** | **0.03** |
| **296_F2 A** | -0.47 | - | -0.36 | 0.62 | - | 0.70 | **0.66** | **0.04** |
| **296_F2 S** | -0.34 | - | -0.38 | 0.71 | - | 0.68 | **0.70** | **0.01** |

[a] A, Albumax; S, Serum

**Supplementary Dataset**

EXCEL datasheet and plots for the relative multiplication rate assays of all the isolates and clones.
